# Visualizing sequential compound fusion and kiss-and-run in live excitable cells

**DOI:** 10.1101/2021.06.21.449230

**Authors:** Lihao Ge, Wonchul Shin, Ling-Gang Wu

## Abstract

Vesicle fusion is assumed to occur at flat membrane of excitable cells. In live neuroendocrine cells, we visualized vesicle fusion at Ω-shape membrane generated by preceding fusion, termed sequential compound fusion, which may be followed by fusion pore closure, termed compound kiss-and-run. These novel fusion modes contribute to vesicle docking, multi-vesicular release, asynchronous release, and endocytosis. We suggest modifying current models of exo-endocytosis to include these new fusion modes.

---

Vesicle fusion releases transmitters, hormones and peptides to mediate many physiological functions, such as synaptic transmission, fight or flight response, and controlling blood glucose level relevant to diabetes^1, 2^. In the last half a century of studies in excitable cells, including neurons and endocrine cells, all models on release steps and modes, such as vesicle docking, fusion pore opening and closure (kiss-and-run), mono- or multi-vesicular release at single release sites, synchronized or asynchronized release, are constructed under a fundamental assumption that vesicles fuse at the flat plasma membrane (PM)^1, 2^. Despite generally accepted, this assumption has not been tested in live cells. Against this concept, sequential compound fusion (Fusion_seq-comp_) – vesicle fusion at a previously fused vesicular Ω-shape structure – has long been proposed in non-excitable cells containing extremely large (~1-5 μm) granules that release contents extremely slowly (~10 s to minutes)^3–5^. Fusion_seq-comp_ could in principle provide a novel mechanism underlying a series of fusion steps and modes, such as vesicle docking, desynchronized multi-vesicular release, asynchronous release, and subsequent endocytosis. These fusion steps and modes may enhance synaptic strength, synaptic reliability, firing information transfer, and the dynamic range of synaptic plasticity and neuromodulation at many synapses^1, 6–8^. However, the concept of Fusion_seq-comp_ remains to be established, because its membrane dynamics, fusion pore, and content release have not been directly visualized and thus proved in any non-excitable or excitable cell. Here we established the Fusion_seq-comp_ concept in excitable cells by direct visualization of its membrane dynamics, fusion pore dynamics and content release dynamics in live cells for the first time.

To visualize membrane dynamics, we transfected EGFP or mNeonGreen attached to phospholipase C delta PH domain (PHG), which binds to PtdIns(4,5)P2 (PIP_2_) at, and thus labels the plasma membrane (PM) in neuroendocrine cells, the primary cultured bovine adrenal chromaffin cell (Fig. 1a)^9, 10^. We added Atto 532 (A532) in the bath, which enters and thus labels fusing vesicles’ Ω-shape profiles (Fig. 1a)^9, 11^. A 1-s depolarization (−80 to +10 mV) via a pipette at the whole-cell configuration induced calcium currents, capacitance changes reflecting exo- and endocytosis, and fusion events observed with stimulated emission depletion (STED) microscopy of PHG/A532 (Fig. 1b-e). Images were acquired at the XZ-plane (near cell bottom) with Y-location fixed at about the cell center for ~1-2 min (XZ/Y_fix_ imaging, 26-300 ms per frame, Fig. 1a); each cell was subjected to only 1 depol1s to avoid whole-cell exo- and endocytosis run-down^9, 11^.

**Figure 1.**
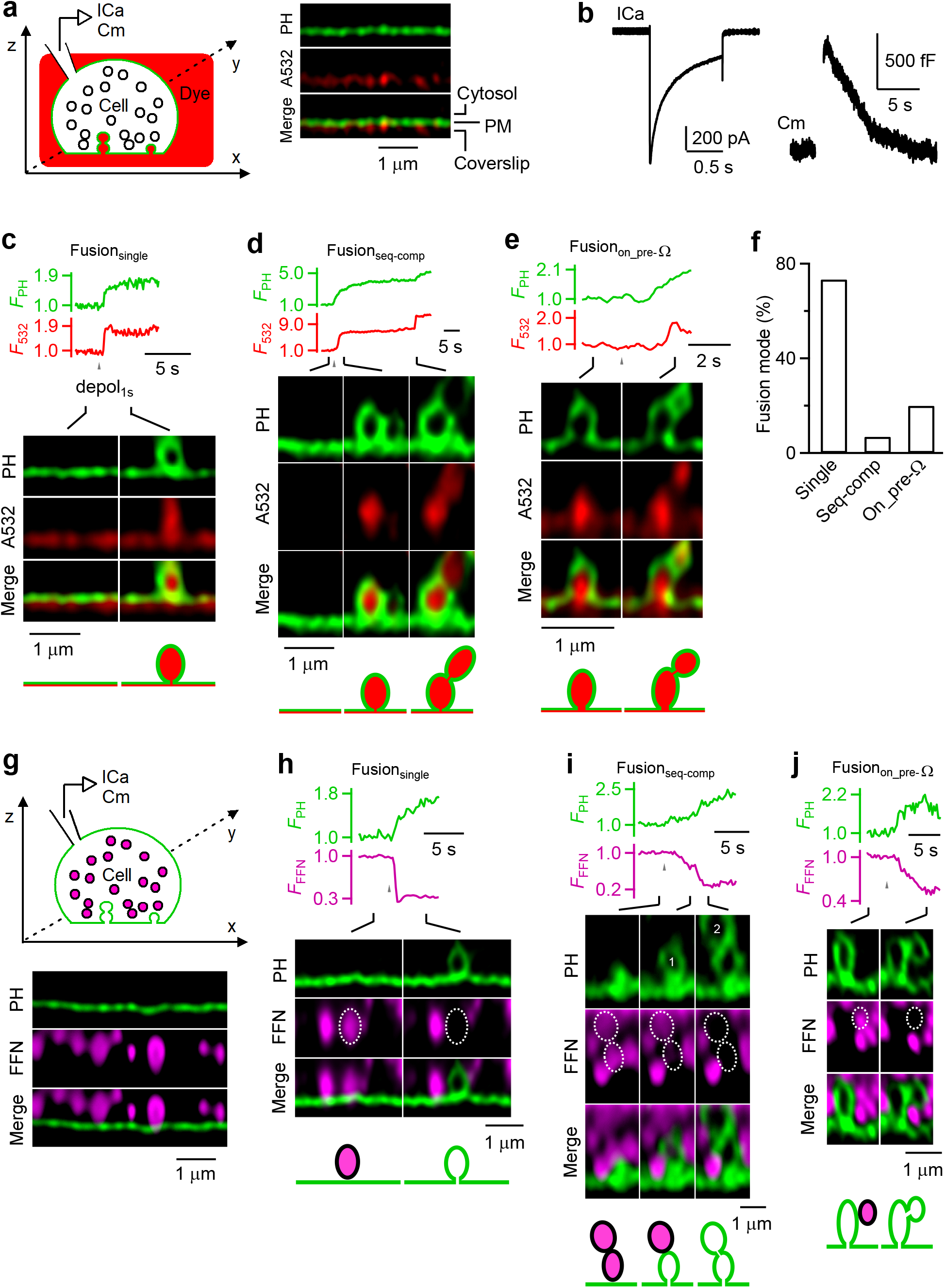
Visualizing sequential compound fusion and fusion on pre-formed Ω-profiles in live cells. **a,** Left: setup drawing. Cell’s membrane is labelled with PH_G_ (green), and bath labelled with A532 (red). ICa and Cm (capacitance) are recorded from the whole-cell pipette. Right: XZ-plane PHG (green) and A532 images for a fraction of a cell (near cell bottom) at rest. Cytosol, PM and coverslip locations are labelled. **b,** Sampled ICa and Cm change induced by depol_1s_. **c-e,** Ω_PH_ fluorescence (F_PH_, normalized to baseline), A532 spot fluorescence (F_532_, normalized to baseline) and STED XZ/Y_fix_ images at times indicated with lines for Fusion_single_ (c), Fusion_seq-comp_ (d), and Fusion_on_pre-Ω_ (e). F_PH_ and F_532_ were collected from fusing vesicle(s). **f,** The percentage of Fusion_single_, Fusion_seq-comp_ and Fusion_on_pre-Ω_ observed with STED XZ/Y_fix_ imaging of PHG/A532 (336 events from 274 cells showing these fusion events). **g,** Similar to panel a, but for imaging FFN511-loaded vesicles (purple, pseudo-colour) and PH_G_-labelled cell membrane (green). **h-j,** F_PH_, FFN511 spot fluorescence (F_FFN_) and STED XZ/Y_fix_ images at times indicated with lines showing release of FFN511 for Fusion_single_ (g), Fusion_seq-comp_ (h, vesicle 1 and 2 are circled and labelled), and Fusion_on_pre-Ω_ (j).

To provide more conclusive proof and to characterize Fusion_seq-comp_, STED PH_G_/A532 imaging data were collected from a large number of cells, 1211 cells, at the voltage-clamp configuration. A total of 336 PH_G_-labelled Ω-shape profiles filled with A532 (Ω_PH_) appeared within a single image frame (26-300 ms), reflecting vesicle fusion that allowed for PH_G_/A532 diffusion from PM/bath into the fusion-generated Ω-profile (Fig. 1c–f). Among 336 Ω_PH_, 247 Ω_PH_ (73.5%) appeared at the flat PM (Fig. 1c, f), reflecting single vesicle fusion (Fusionsingle, for detail, see Ref. ^9, 10, 12^); 23 Ω_PH_ (6.9%) appeared at flat PM, but followed at 0.2-85 s later on its top by a sudden appearance of another Ω_PH_, forming an 8-shape structure reflecting Fusion_seq-comp_ (Fig. 1d, f); 66 Ω_PH_ (19.6%) appeared on the top of Ω_PH_ preformed before depol_1s_ (Fusion_on_pre-Ω_), which also formed an 8-shape structure (Fig. 1e, f). Preformed Ω_PH_ could be from previous fusion events that maintained a Ω-shape, as recently reported^9, 11^. Supporting this possibility, the 2^nd^ fusion may occur ~20-85 s after 1^st^ fusion during Fusion_seq-comp_ (e.g., Fig. S1, n = 4). Thus, Fusion_on_pre-Ω_ may reflect Fusion_seq-comp_ with a prolonged delay.

To demonstrate the release dynamics of Fusion_seq-comp_ and Fusion_on_pre-Ω_, we loaded vesicles with fluorescent false neurotransmitter FFN511, a substrate for vesicle monoamine transporter, via bath application (Fig. 1g)^13^. XZ/Y_fix_ imaging of PH_G_/FFN511 revealed decrease of FFN511 spot fluorescence (F_FFN_) and simultaneous appearance of Ω_PH_ at the same spot, reflecting fusion-generated Ω_PH_ that releases FFN511 (Fig. 1h–j). FFN511 releasing spots may 1) fuse on flat PM, reflecting Fusion_single_ (Fig. 1h, n = 153), 2) fuse on flat PM, but followed on its top by the 2^nd^ fusion that released its FFN511 content and created the 2^nd^ Ω_PH_, (forming an 8-shape structure with the 1^st^ Ω_PH_), reflecting Fusion_seq-comp_ (Fig. 1i, n = 11), or 3) fuse on preformed Ω_PH_ to form a PH_G_-labelled 8-shape structure, reflecting Fusion_on_pre-Ω_ (n = 31, Fig. 1j). These results established the concept of Fusion_seq-comp_ and Fusion_on_pre-Ω_ by demonstrating their vesicular positions and content release.

Next, we examined fusion pore and Ω_PH_ membrane dynamics of Fusion_seq-comp_ and Fusion_on_pre-Ω_. As previously characterized^9, 10^, Ω_PH_ in Fusion_single_ may maintain an open pore (stay-fusion, Fig. 2a), close its pore at ~0.05 - 30 s later (close-fusion, Fig. 2b), or shrink to merge with the plasma membrane (shrink-fusion, Fig. 2c; summarized in Fig. 2d). Close-fusion was detected as A532 fluorescence (F_532_, strongly excited) dimming due to pore closure that prevented bath fluorescent A532 from exchanging with bleached A532, while PHG fluorescence (FPH, weakly excited) sustained or decayed with a delay that reflected PtdIns(4,5)P_2_ conversion into PtdIns(4)P and/or vesicle pinch off (Fig. 2b); stay-fusion was detected as sustained F_532_ and F_PH_ (Fig. 2a); shrink-fusion, Ω_PH_ shrinking with parallel decreases of F_532_ and F_PH_ (Fig. 2c) (for detail, see Refs. ^9, 10^).

**Figure 2.**
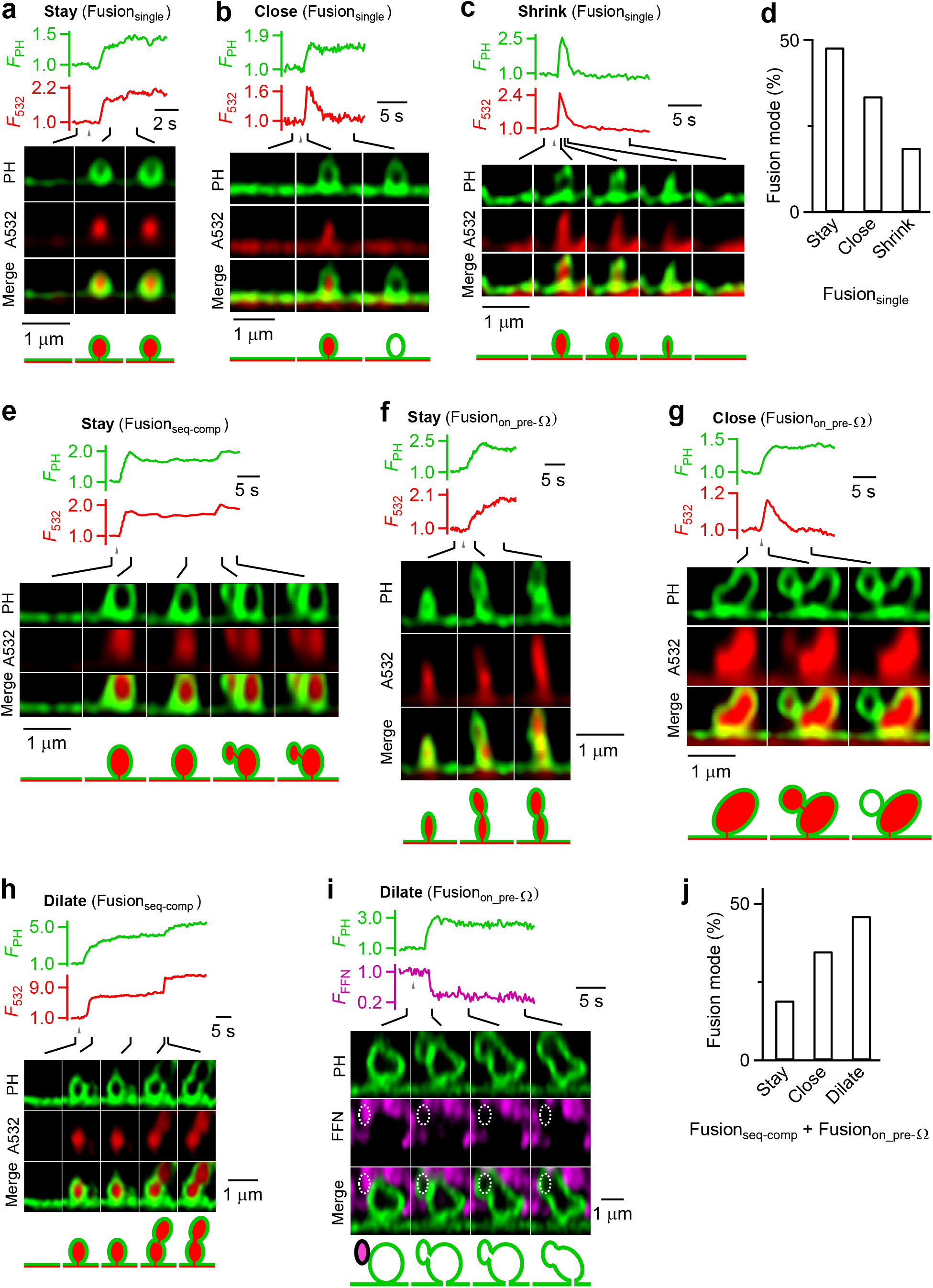
Pore dynamics of sequential compound fusion and fusion on pre-formed Ω-profiles. **a-c,** F_PH_, F_532_, and sampled images for three modes of Fusion_single_: stay-fusion (a), close-fusion (b), and shrink-fusion (c). **d,** Stay-, close- and shrink-fusion percentage for Fusionsingle (247 events, 212 cells). **e-f,** FPH, F532, and sampled images showing stay-fusion for Fusion_seq-comp_ (e, stay-fusion refers to 2nd fusion event) and for Fusion_on_pre-Ω_ (f). **g,** F_PH_, F_532_, and sampled images showing close-fusion for Fusion_on_pre-Ω_. **h,** F_PH_, F_532_, and sampled images showing dilation of the fusion pore during Fusion_seq-comp_ (pore dilation refers to 2nd vesicle fusion) **i,** F_PH_, F_511_, and sampled images showing dilation of the fusion pore during Fusion_on_pre-Ω_. **j,** Percentage of stay-fusion, close-fusion and pore dilation for the vesicle that fused at the previously generated Ω_PH_ (n = 89). Data from Fusion_seq-comp_ (n = 23) and Fusion_on_pre-Ω_ (n = 66) were pooled together.

The 2^nd^ Ω_PH_ in Fusion_seq-comp_ and Fusion_on_pre-Ω_ may 1) remain unchanged with an open pore, reflected as sustained F_532_ and F_PH_ (Fig. 2e, f), analogous to Fusionsingle’s stay-fusion (Fig. 2a), 2) close its pore at ~0.05-30 s later, reflected as F_532_ decay while F_PH_ sustained or decayed with a delay (Fig. 2g), analogous to Fusion_single’s_ close-fusion (Fig. 2b), or 3) dilate its pore till the 8-shape was converted to an elongated or large Ω-shape (e.g., Fig. 2h, i; Fig. 2j shows their percentages). We termed 2^nd^ Ω_PH_ close-fusion (e.g., Fig. 2g) compound kiss-and-run, a new form of kiss-and-run. Unlike Fusion_single_, we did not observe 2^nd^ Ω_PH_ shrink-fusion, but pore dilation (Fig. 2h–j).

The 20-80% FFN511 fluorescence decay time was similar between the 2^nd^ and the 1^st^ Ω_PH_ during Fusion_seq-comp_ (Fig. 3a, b), suggesting that Fusion_seq-comp_ releases vesicular contents as efficiently as Fusion_single_. However, the 2^nd^ Ω_PH_ appeared at ~0.2-85 s after the 1^st^ Ω_PH_ during Fusion_seq-comp_ (e.g., Figs. 1d, 1i, 2e; summarized in Fig. 3c), indicating that Fusion_seq-comp_ can generate desynchronized multi-vesicle release at single release sites. Given that the 1^st^ Ω_PH_ occurred mostly during and within ~1 s after depol1s (Fig. 3c), the 1^st^ Ω_PH_ reflected synchronized release; the various time delay of the 2^nd^ Ω_PH_ (Fig. 3c) thus reflected asynchronous release. We concluded that Fusion_seq-comp_ contributes to the generation of desynchronized multi-vesicular release and asynchronous release.

**Figure 3.**
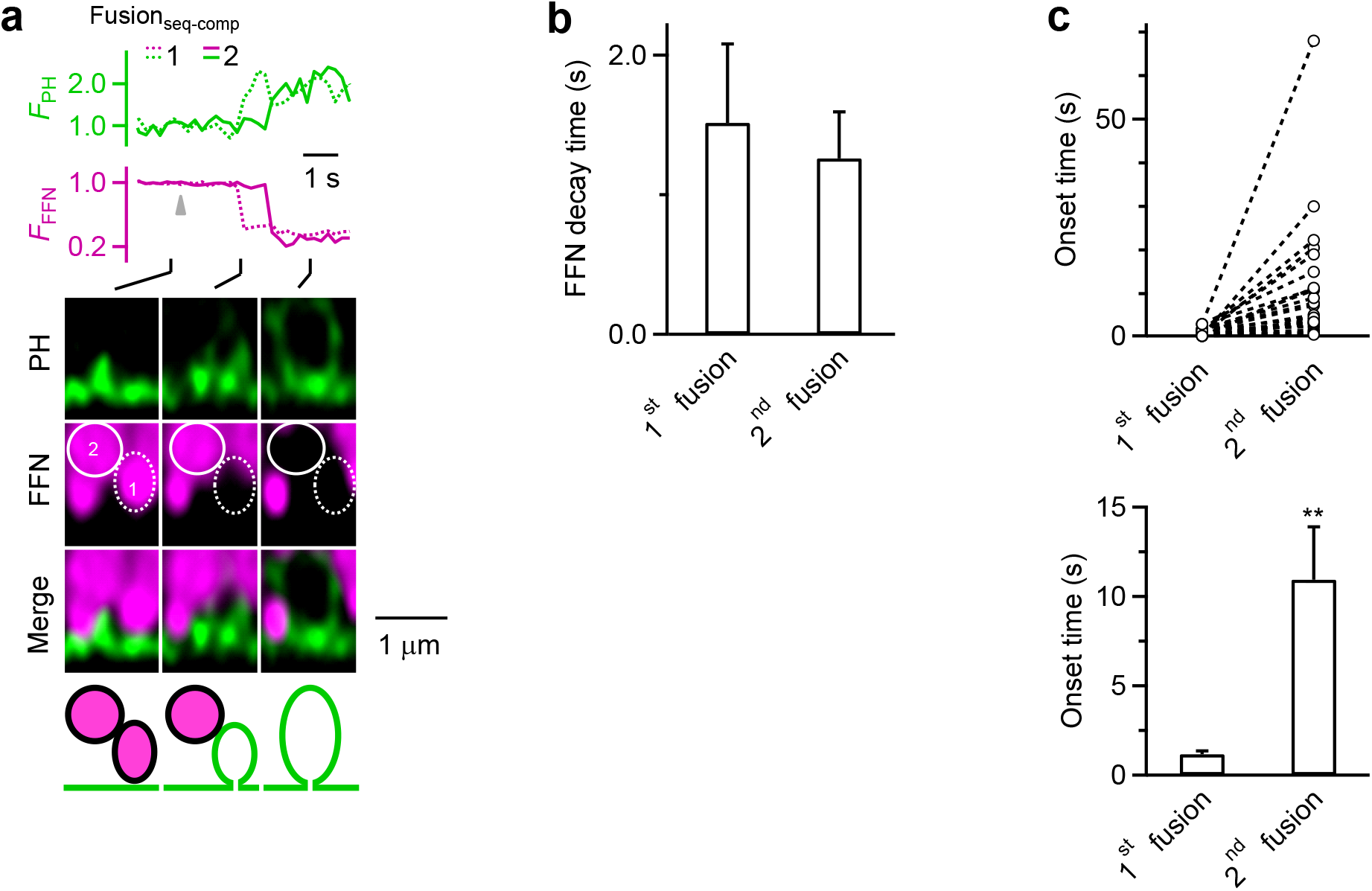
Sequential compound fusion releases contents efficiently, but generates desynchronized multi-vesicular release and asynchronized release at single release sites. **a,** F_PH_, F_FFN_, and sampled images of the 1^st^ (dotted circle, dotted trace) and the 2^nd^ (solid circle, solid trace) fusion for a Fusion_seq-comp_ event. **b,** The 20-80% F_FFN_ decay (release) time (mean + s.e.m.) for 1^st^ and 2^nd^ fusion during Fusion_seq-comp_ (11 events, 11 cells under PHG/FFN511 imaging). No significant difference was found (paired t test, p = 0.712). **c,** Upper: the onset time of the 1^st^ and the 2^nd^ fusion during Fusion_seq-comp_ (23 events, 23 cells under PHG/A532 imaging). 1 fusion per circle; dash lines connect two fusion events from the same Fusion_seq-comp_. Onset time 0 refers the onset of depol1s. Lower: mean onset (+ s.e.m., n = 23) of the 1^st^ and the 2^nd^ fusion during Fusion_seq-comp_ (**: p < 0.01, paired t-test). Left and right panels are from the same data set.

The present work firmly established the concept of sequential compound fusion and compound kiss-and-run by directly visualizing their membrane, fusion pore and content-release dynamics in live cells for the first time. Establishing these new concepts in excitable cells may conceptually advance our understanding of secretory vesicle exo-endocytosis, because sequential compound fusion and compound kiss-and-run may constitute new mechanisms contributing to the generation of desynchronized multi-vesicular release at single release sites, asynchronous release, vesicle docking and priming, and vesicle endocytosis, as discussed below.

Multi-vesicular release from single release sites enhance synaptic strength, synaptic reliability, and the dynamic range of synaptic plasticity and neuromodulation at many synapses^1^. The mechanism underlying multi-vesicular release at single release sites is poorly understood^1^. Sequential compound fusion readily explains how single release sites produce multi-vesicular release, particularly the desynchronized multi-vesicular release, which enhances the precise and efficient firing information transfer at synapses^6, 7^. If the release interval of sequential compound fusion is minimal, it may also explain the coordinated or simultaneous multi-vesicular release at single release sites^1, 14^.

While depolarization-evoked release is mostly synchronized, asynchronous release lasting much longer than the brief depolarization also takes place in many excitable cells, which may transfer a brief presynaptic firing burst into a prolonged postsynaptic firing burst at synapses^8^. The differences in the mechanisms underlying asynchronous and synchronous release remain not well understood. Different calcium sensors with different calcium affinity have been suggested^8^. Sequential compound fusion evidently generates a delay in releasing the second vesicle (Fig. 3), providing a novel mechanism contributing to the generation of asynchronous release.

Vesicle fusion must involve vesicle movement towards the PM release sites for docking and subsequent vesicular V-SNARE and PM T-SNARE binding that may prime docked vesicles for release^8, 15^. Our findings suggest modifying this concept by including a new mechanism of generating release sites for docking – the fusion-generated Ω-profiles are maintained and serve as the new release sites for docking and priming. Such a docking process saves vesicles from spending energy to travel one-vesicle-length of distance for docking at the flat PM. The priming process may involve diffusion of T-SNARE from the PM to the fusion-generated vesicular Ω-profile and T-SNARE binding with V-SNARE of the docking vesicle^16^. These processes may take more time^16^, explaining the prolonged release interval of sequential compound fusion that produces asynchronous release.

Compound kiss-and-run reported here is a new mode of exo-endocytosis that retrieves vesicles undergoing sequential compound fusion. It may explain the electron microscopic observation of 8-shape (or sausage-shape) structures and large vesicles that are otherwise interpreted as different mechanisms, such as vesicle budding, bulk endocytosis of large vesicles, and/or cytosol vesicle-vesicle fusion. We suggest modifying current models of secretory vesicle endocytosis^2, 17^ by including compound kiss-and-run as a new mode of endocytosis.

While obtained from chromaffin cells containing large dense-core vesicles, our findings are most likely applicable to neurons for two reasons. First, like neurons, chromaffin cells are excitable cells with a neuronal origin and very similar calcium-, synaptotagmin-, SNARE-, and dynamin-dependent exo- and endocytosis^2, 18^. Second, neuron contains large dense-core and small clear-core vesicles, both of which may maintain an Ω-shape after fusion^19–22^, the prerequisite for mediating sequential compound fusion. Direct visualization is needed to prove sequential compound fusion and kiss-and-run in neurons. Visualizing sequential compound fusion of small synaptic vesicles (~30-50 nm) requires much higher spatial-temporal resolution than what we have (~60 nm/26-300 ms). Significant technical advancements are needed to overcome this technical problem in the future.

Cytosolic vesicle-vesicle fusion is the first form of compound fusion being proposed, based on the observation of 1) cytosolic 8-shape or sausage-like vesicular structures, and 2) capacitance jumps and synaptic miniature currents (quantal size) too large for single vesicle fusion in non-excitable and excitable cells, such as pancreatic acinar cells, eosinophils, mast cells, calyx-type and ribbon-type synapses^3, 5, 23–27^. However, direct observation of the membrane transformation during vesicle-vesicle fusion, which can fully establish the concept of vesicle-vesicle fusion is still missing. Similar to vesicle-vesicle fusion, sequential compound fusion was suggested in non-excitable cells based on the observation of 1) sequential release of lysotracker green loaded into the extremely large (~1-5 μm) granules in eosinophils^3^, and 2) sequential generation of extracellular-dye-loaded extremely large tube-like structures from the PM into the cytosol in acinar cells^4^. However, these observations could not fully exclude the possibility that the extremely large, apparently cytosolic structure could be docked at PM that was not visible at the imaged plane, or that the extracellular-dye-loaded structure reflects endocytic membrane invagination (content release is not imaged simultaneously). Because of these uncertainties, and most importantly the lack of evidence showing direct, simultaneous membrane, pore and release dynamics of sequential compound fusion in live cells, the concept of sequential compound fusion is not fully established^2^. The present work provided the missing evidence required to fully establish the concept of sequential compound fusion – the dynamics of membrane transformation, fusion pore, and content release. Furthermore, we link this concept to a new endocytic mode, compound kiss-and-run, and extend these concepts to excitable cells that release much smaller vesicles rapidly. Thus, sequential compound fusion and compound kiss-and-run may be a widespread exo-endocytosis mode used by excitable and non-excitable cells to release vesicular contents that may mediate important functions such as neuronal communication, fight or flight response, regulation of blood glucose level relevant to diabetes, and immune responses^1, 2^. The technique we used here opens the door to study the functions and mechanisms of sequential compound fusion in live cells.

## Acknowledgements

This work was supported by the National Institute of Neurological Disorders and Stroke Intramural Research Program (ZIA NS003009-15 and ZIA NS003105-10) to L.G.W. We thank Carolyn Smith for STED microscopy support.

## Author Contributions

L.G. did most FFN511-related experiments. W.S. did most PHG/A532-related experiments. L.G.W. supervised the project. L.G. wrote the experimental results; L.G.W. wrote the manuscript with help from L.G. and W.S.

## Materials and Methods

### Chromaffin cell culture

We prepared primary bovine adrenal chromaffin cell culture as described previously^11^. Fresh adult (21 - 27 months old) bovine adrenal glands (from a local abattoir) were immersed in pre-chilled Locke’s buffer on ice containing: NaCl, 145 mM; KCl, 5.4 mM; Na2HPO4, 2.2 mM; NaH2PO4, 0.9 mM; glucose, 5.6 mM; HEPES, 10 mM (pH 7.3, adjusted with NaOH). Glands were perfused with Locke’s buffer, then infused with Locke’s buffer containing collagenase P (1.5 mg/ml, Roche), trypsin inhibitor (0.325 mg/ml, Sigma) and bovine serum albumin (5 mg/ml, Sigma), and incubated at 37°C for 20 min. The digested medulla was minced in Locke’s buffer, and filtered through a 100 μm nylon mesh. The filtrate was centrifuged (48 xg, 5 min), re-suspended in Locke’s buffer and re-centrifuged until the supernatant was clear. The final cell pellet was re-suspended in pre-warmed DMEM medium (Gibco) supplemented with 10% fetal bovine serum (Gibco).

### Electroporation and plating

Cells were transfected by electroporation using Basic Primary Neurons Nucleofector Kit (Lonza), according to the manufacturer’s protocol and plated onto glass coverslips with mouse Laminin coating over PDL layer (Neuvitro). The cells were incubated at 37°C with 9% CO2 and used within 5 days.

### Plasmids and fluorescent dyes

The PH-EGFP (phospholipase C delta PH domain attached with EGFP) was obtained from Dr. Tamas Balla. PH-mNeonGreen construct was created by replacing the EGFP tag of PH-EGFP with mNeonGreen (Allele Biotechnology) ^28^. Both PH-EGFP and PH-mNeonGreen are abbreviated as PHG. For Atto 532 (A532, Sigma) imaging, A532 concentration in the bath solution was 30 μM. For FFN511 (Abcam) imaging, cells were bathed with FFN511 (5-10 μM) for 10 min and images were performed after washing out FFN511 in the bath solution.

Overexpression of PHG did not significantly affect the basic properties of exo- and endocytosis, because 1) whole-cell capacitance measurements and imaging show robust exo- and endocytosis, and similar percentages of close-fusion and non-close-fusion as control^10, 11^, and 2) PHG-labelled fusion pore could also be observed with imaging of extracellularly applied mCLING-A488 or with EM^9^.

### Electrophysiology

At room temperature (20 - 22°C), whole-cell voltage-clamp and capacitance recordings were performed with an EPC-10 amplifier together with the software lock-in amplifier (PULSE, HEKA, Lambrecht, Germany) ^11, 29^. The holding potential was -80 mV. For capacitance measurements, the frequency of the sinusoidal stimulus was 1000 - 1500 Hz with a peak-to-peak voltage ≤ 50 mV. The bath solution contained 125 mM NaCl, 10 mM glucose, 10 mM HEPES, 5 mM CaCl2, 1 mM MgCl2, 4.5 mM KCl, and 20 mM TEA, pH 7.3 adjusted with NaOH. The pipette (2 – 4 MΩ) solution contained 130 mM Cs-glutamate, 0.5 mM Cs-EGTA, 12 mM NaCl, 30 mM HEPES, 1 mM MgCl2, 2 mM ATP, and 0.5 mM GTP, pH 7.2 adjusted with CsOH. These solutions pharmacologically isolated calcium currents.

For stimulation, we used a 1-s depolarization from the holding potential of -80 mV to +10 mV (depol1s). We used this stimulus, because it induces robust exo-endocytosis as reflected in capacitance recordings (Fig. 1a) ^11, 30, 31^. In a fraction of experiments during FFN511 imaging, we used 10 pulses of 400-ms depolarization from -80 mV to +10 mV at 2 Hz, which evoked more fusion events.

### STED imaging

STED images were acquired with Leica TCS SP8 STED 3× microscope that is equipped with a 100 x 1.4 NA HC PL APO CS2 oil immersion objective and operated with the LAS-X imaging software. Excitation was with a tunable white light laser and emission was detected with hybrid detectors. In time-gated STED mode, PH-EGFP and A532 were sequentially excited at 470 and 532 nm, respectively, with the 592 nm STED depletion beam, and their fluorescence collected at 475-525 nm and 540-587 nm, respectively. PH-mNeonGreen and A532 were sequentially excited at 485 and 540 nm, respectively, with the 592 nm STED depletion beam, and their fluorescence collected at 490-530 nm and 545-587 nm, respectively. PH-mNeonGreen and FFN511 were sequentially excited at 505 and 442 nm, respectively, with the 592 nm STED depletion beam, and their fluorescence collected at 510-587 nm and 447-490 nm, respectively.

The excitation power for A532 was 10% of the maximum, at which fluorescent A532 can be bleached within a few seconds. This feature was used to distinguish whether the fusion pore is closed or not, because pore closure prevents bleached A532 (caused by strong excitation) from exchange with fluorescent A532 in the bath, resulting in A532 spot fluorescence decay^9-11^. In contrast, an open pore would not cause A532 spot fluorescence decay, because an open pore allows for continuous exchange of bleached A532 in the Ω-profile with fluorescent A532 in the bath ^9–11^.

STED imaging generally causes more photobleaching and phototoxicity. Severe phototoxicity could cause loss of the whole-cell giga seal during patch-clamp recording ^11^. In general, we avoided severe phototoxicity by applying only one depol1s and imaging for ~1-2 minper cell. With this setting, we have not noticed significant differences in the exo- and endocytosis properties obtained under confocal and STED imaging conditions^10, 11^. For imaging of PHG and A532, continuous exchange of bleached PHG or A532 with fluorescent ones from non-imaging areas lessened the photobleaching problem.

### STED scanning modes

STED images were acquired at the cell bottom at the XZ-plane (perpendicular to the coverslip) with the Y-axis location fixed at about the cell center (Figure 1a, XZ/Y_fix_ scanning mode). We repeated XZ/Y_fix_ scanning every 26-300 ms at 15 nm per pixel in an XZ area of 19.4 μm x 0.7-1.5 μm^9^.

The STED resolution for imaging PHG (PH-EGFP or PH-mNeonGreen) in our conditions was ~60 nm on the microscopic X- and Y-axis (parallel to cell-bottom membrane or coverslip), and ~150-200 nm on the microscopic Z-axis. STED images were deconvolved using Huygens software (Scientific Volume Imaging).

### Data selection

For every cell recorded with a pipette under the whole-cell configuration, the data within the first 2 min at the whole-cell configuration were used, which avoided rundown of endocytosis (gradual disappearance of endocytosis) as previously reported under the whole-cell configuration for a long time^11, 32^. Cells expressed with PHG were used for visualization of fusion events. The criteria for selecting PHG-labelled Ω for analysis during XZ scanning are described in Figure S2 of Shin et al., 2018.

### Analysis of PH_G_-labelled Ω-shape profiles

STED images of PH-Ω were analyzed with ImageJ and LAS X (Leica). During XZ scanning, some depol1s-induced PH-Ω-profiles were out of the same Y focal plane, as the outline of the Ω-profile was vague or unclear (for detail, see ^9^). These out-of-focus Ω-profiles were not included for analysis.

Pores labelled with PHG were identified based on the image and the fluorescence intensity line profile (for detail, see ^9^). We first identified the fluorescently labelled Ω-profiles with an open pore, the edge of which was continuous with PM. The intensity line profile in the pore region should show a valley with a peak at least three times larger than the baseline fluctuation (standard deviation) in the non-pore region (for detail, see ^9^). The full-width-half-maximum of the valley of the intensity line profile across the pore was proportional to the pore diameter, as shown with simulation^9^. Pore dilation of the fusing Ω_PH_ during Fusion_seq-comp_ or Fsuion_pre-Ω_ was judged with eyes.

Identification of stay-, close- and shrink-fusion during XZ/Y_fix_ STED imaging of PH_G_/A532 were described in detail previously^9^. During XZ/Y_fix_ imaging, A532 was excited at a high laser power so that fluorescent A532 can be bleached with a time constant of 1.5-3.5 s. Pore closure was identified as the gradual dimming of the A532 spot fluorescence to baseline during XZ PHG/A532 imaging while PHG image remained unchanged or dimmed gradually without changing the Ω_PH_ size^9^. A532 fluorescence dimming was not due to a narrow pore smaller than A532 molecule size, because after A532 spot dimming, bath application of an acid solution cannot quench the pH-sensitive VAMP2-EGFP or VAMP2-pHluorin overexpressed at the same spot, indicating that the spot is impermeable to H^+^ or OH^−^, the smallest molecules, and thus is closed^10, 11^.

### Statistical tests

Data were expressed as mean ± s.e.m. Replicates are indicated in results and figure legends. N represents the number of cells, fusion events, pores, or Ω-profiles as indicated in results and figure legends. The statistical test used is *t* test or ANOVA. Although the statistics were performed based on the number of cells, fusion events, pores, and Ω-profiles, each group of data was collected from at least four primary chromaffin cell cultures. Each culture was from at least two glands from one bovine.

