## Supplemental file for "Visualizing sequential compound fusion and kiss-and-run in live excitable cells"

### Supplementary Information

Supplementary information contains Figure S1.

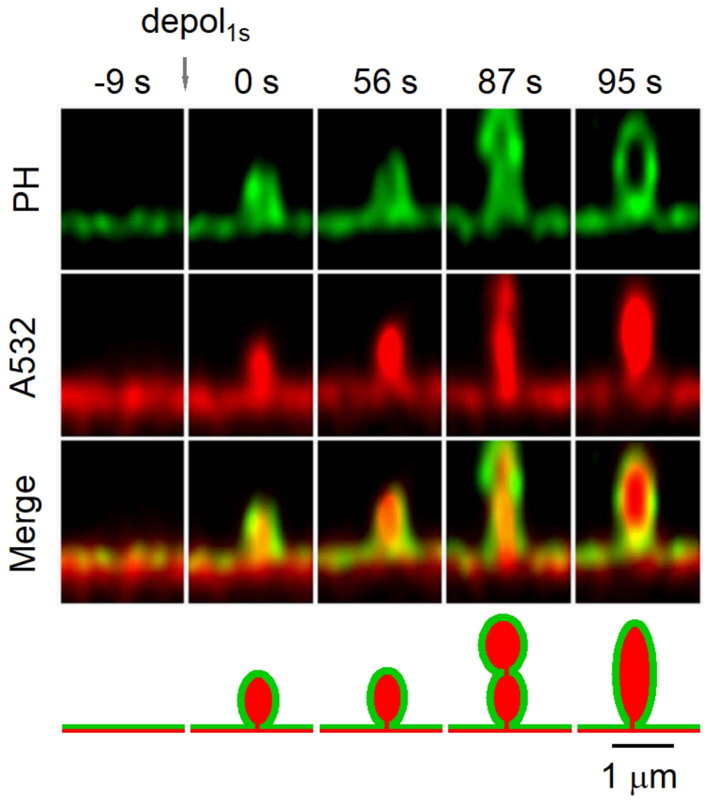


**Figure S1. A sequential compound fusion with a prolonged interval between the 1st and the 2nd fusion**

PHG image, A532 image, and their merge images at various times during a sequential compound fusion event. The time is labelled relative to the onset of the 1st fusion (0 s). The 2nd fusion occurred at 85 s later. The schematic drawing is also shown at the bottom.
